## Supplemental Figure 1 for "SMAD4 promotes somatic-germline contact during murine oocyte growth"

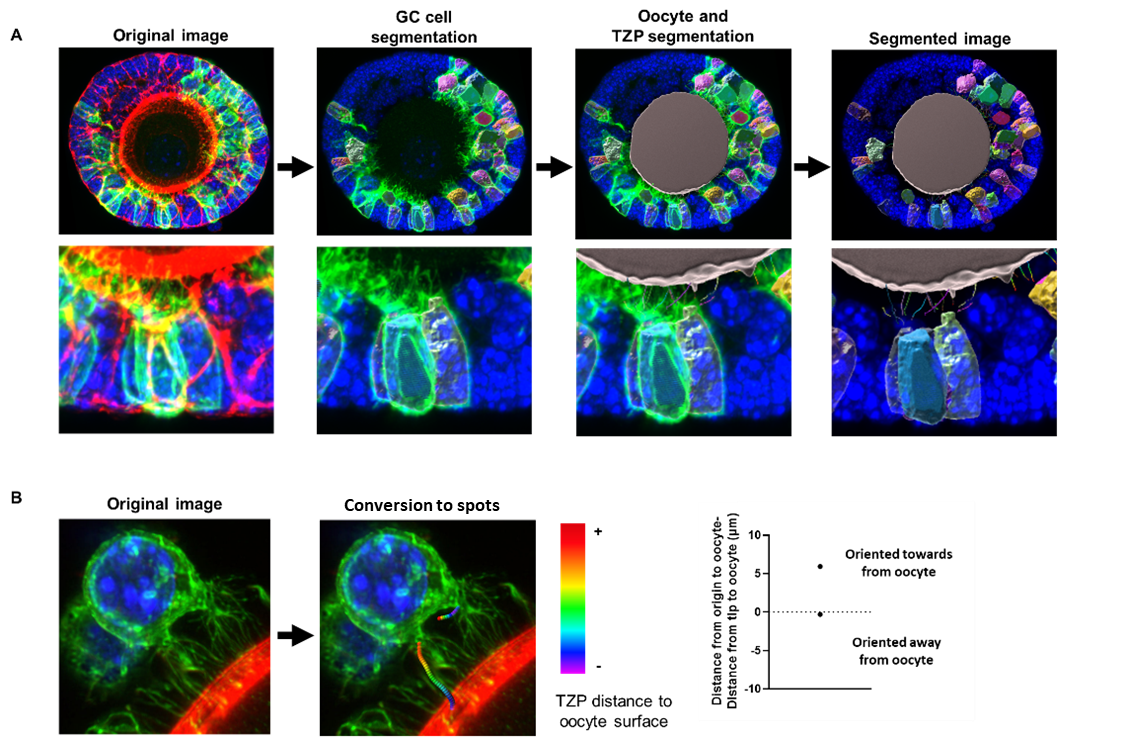

**Supplemental Figure 1. Segmentation pipeline of GFP-positive granulosa cells.** (A) 3D reconstruction of granulosa cells, oocyte and TZPs. (B) TZP segmentation into spots. Graph shows an example of spot subtraction of two TZPs oriented towards and away from the oocyte. Heatmap depicts the gradient of colours for the spots that are closer (violet) and farther from the oocyte (red).
